# Helping Behavior in Prairie Voles: A Model of Empathy and the Importance of Oxytocin

**DOI:** 10.1101/2020.10.20.347872

**Authors:** Kota Kitano, Atsuhito Yamagishi, Kengo Horie, Katsuhiko Nishimori, Nobuya Sato

## Abstract

Several studies suggest that rodents show empathic responses and helping behavior toward others. We examined whether prairie voles (*Microtus ochrogaster*) would help conspecifics who were soaked in water by opening a door to a safe area. Door-opening latency decreased as task sessions progressed. When the conspecific was not soaked in water, the latency of the door-opening did not decrease, indicating that the distress of the conspecific is necessary for the learning of the door-opening. These suggest that prairie voles learn helping behavior. We found no sex difference in the door-opening latency or the interest in other voles soaked in water. Additionally, we also examined the helping behavior in prairie voles in which oxytocin receptors were genetically knocked out. Oxytocin receptor knockout voles demonstrated less learning of the door-opening and less interest in other voles soaked in water. This suggests that oxytocin is important for the emergence of helping behavior.

## INTRODUCTION

Empathy is an innate ability to experience and share the mental state of others (Decety et al., 2016; de Waal and Preston, 2017; Meyza et al., 2017; Preston and de Waal, 2002). In mammals, prosocial behaviors such as helping and consolation are essential for the development of society and are thought to be elicited by empathy (de Waal and Preston, 2017). The existence of empathy has been suggested in a variety of species, including non-human primates (Campbell and de Waal, 2011; Koski and Sterck, 2010; Pruetz, 2011), dogs (Palagi et al., 2015), birds (Gallup et al., 2015), and even rodents (Bartal et al., 2011; Bartal et al., 2016; Sato et al., 2015). Several studies have examined empathy in rodents using behavioral indicators, such as vicarious learning of fear (Atsak et al., 2011; Pisansky et al., 2017) and helping behavior like releasing others from a distressed situation (Bartal et al., 2011; Bartal et al., 2016; Sato et al., 2015; Yamagishi et al., 2019; Yamagishi et al., 2020). However, understanding of empathy in rodents is still inadequate. It has been shown that helping behavior, in which an actor pays a cost and gives a benefit to others, is based on empathy. Helping behavior has been observed in animals that are considered to be highly intelligent, such as chimpanzees (Pruetz, 2011; Yamamoto et al., 2012), elephants (Schulte, 2000), and dolphins (Kuczaj et al., 2015). Recent studies suggest that rats also demonstrate helping behavior toward distressed conspecifics, such as in a narrow tube (Bartal et al., 2011; Bartal et al., 2016) and in water (Sato et al., 2015; Yamagishi et al., 2019; Yamagishi et al., 2020). Detecting others’ distress is a prerequisite for helping behavior (Cronin, 2012; Decety et al., 2016). This suggests that rats may have an ability of social cognition that allows them to individually recognize conspecifics and one of empathy that allows them to perceive the distress of others.

Prairie voles (*Microtus ochrogaster*) are known to show more of various social behaviors, such as social bonding, nurturing, allogrooming, and huddling, compared to other rodents such as mice and rats (Aragona and Wang, 2004; Carter and Getz, 1993; Getz and Carter, 1996; Getz and Hofmann, 1986). Because of this feature, many studies on social behavior have been carried out using prairie voles (Aragona and Wang, 2004; Young and Wang, 2004; Tabbaa et al., 2017). Previous studies suggest that prairie voles show empathy-like behavior, such as freezing caused by emotional contagion of conspecifics’ fear, and consolation (Burkett et al., 2016; Stetzik et al, 2018; Wardwell et al., 2020). However, helping behavior that rescues conspecifics from distressed situations has not been investigated in prairie voles. The purpose of this study was to investigate whether prairie voles display helping behavior using a door-opening paradigm, which has been used to examine helping behavior in rats (Bartal et al., 2011; Sato et al., 2015).

Several studies have reported that oxytocin and oxytocin receptors in rodents affect sociality, including affiliative behavior, social cognition, and empathic response (Bartz et al., 2010; Marlin and Froemke, 2017; Rogers-Carter et al., 2018; Ross and Young, 2009; Winslow and Insel, 2002; Young and Barrett, 2015). The role of oxytocin in helping behavior is largely unexamined except in studies that examined the effect of oxytocin administration and of blocking of oxytocin receptors in rats (Yamagishi et al., 2020; Yamagishi et al., 2019). In this study, we examined the effects of oxytocin on helping behavior using oxytocin receptor knockout (OXTrKO) voles.

## RESULTS

### Prairie Voles Show Helping Behavior Quickly

Fourteen pairs of same-sex prairie voles (seven pairs of males and seven pairs of females) were tested with a door-opening task. One member of each pair was assigned to be a soaked vole, and the other was assigned to be a helper vole. The experimental apparatus consisted of a pool area and a ground area (Figures 1A and 1B). The helper vole in the ground area rescued the soaked vole in the pool area by opening a circular door (see Supplemental Video 1). The helper vole could see the soaked vole through a transparent acrylic plate to which the circular door was attached. The task session started when the helper vole was placed in the ground area after the soaked vole was placed in the pool area. We measured the latency of door-opening. After the helper vole opened the door within 10 minutes, the helper and soaked voles were allowed to interact for two minutes. Before the door-opening task, the helper voles were placed in the ground area for five minutes for two days to habituate to the experimental apparatus. Because the circular door was opened and placed on the floor of the ground area during the habituation, the helper voles could go back and forth between the ground and pool areas. We measured the helper voles’ preference for water by counting the time they spent in the ground and the pool areas.

**Figure 1.**
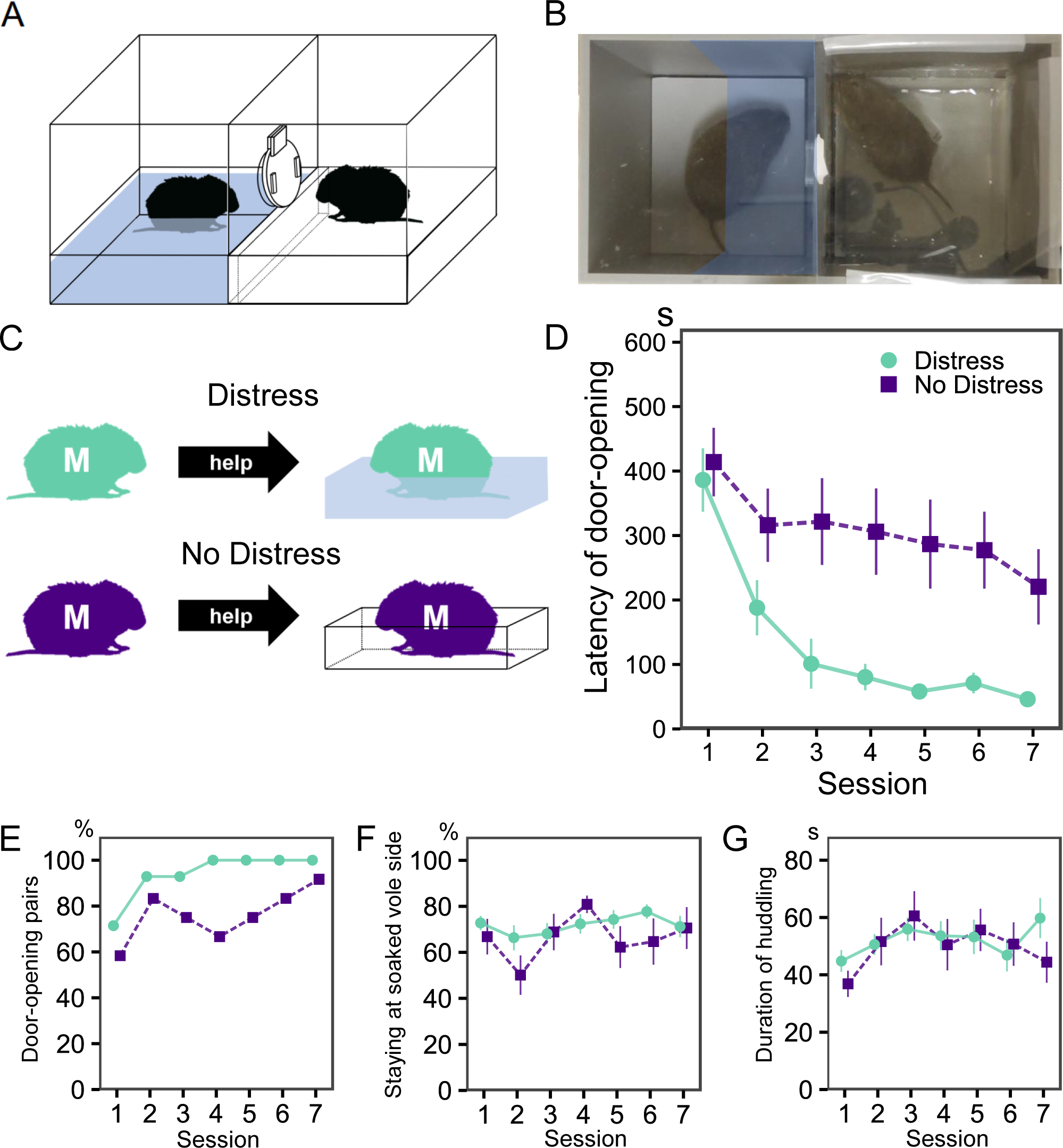
Experimental apparatus and behavioral results in wildtype voles. (A) Schematic diagram of the experimental apparatus. A helper vole could rescue a soaked vole from the distressed situation of being in water by opening the circular door. (B) A photo image during the door-opening task. An acrylic plate covered the ground and pool areas, which reflected the video camera and the ceiling of the laboratory. The blue region is defined as the soaked vole side. (C) There were two groups of paired voles. In one group, one of the pair (a cagemate of the helper vole) was soaked in water (Distress). In the other group, the cagemate was not soaked in water (No Distress). The mean latency (D) and the percentage (E) of door-opening in the helper voles of the Distress and No Distress groups. (F) The percentage of time that the helper voles stayed on the soaked vole side in the Distress and No Distress groups (The data of three voles with incomplete video recordings were excluded). (G) The mean duration of huddling after opening the door in the Distress and No Distress groups. The error bars indicate the standard error of means.

The latency of the door-opening decreased over the seven sessions (Figure 1D and Figure S1). All of the helper voles showed door-opening behavior by the fourth session (Figure 1E). A one-way analysis of variance (ANOVA) revealed a significant effect of session (*F*(6, 78) = 18.34, *p* < .001). This suggests that the helper voles learned the door-opening behavior to free the soaked voles from the pool area. During the habituation period, the helper voles stayed longer in the ground area (550.50±6.54 s, mean±SD) than the pool area. The time spent in the ground area was significantly longer than the expected value (300 s, half of the habituation period, *t*(13) = 36.93, *p* < .001). This suggests that prairie voles have an aversion to water, and that the helper voles rescue the soaked voles from distress by opening the circular door.

The door-opening behavior was potentially learned through factors other than empathy, for instance, social interaction. To examine the possibility that helping behavior is motivated by empathy, we tested the door-opening behavior of the helper voles when the cagemate was not soaked in water (Figure 1C; see Supplemental Video 2). The helper voles (n = 12; all of them were male), which were different from those used in the previous experiment, did not demonstrate a substantial decrease in the latency of door-opening when their cagemate was in the ground area instead of soaked in water (Figure 1D and Figure S2). A one-way ANOVA revealed no effect of session (*F*(6, 66) = 1.59, p = .163). We compared the door-opening latencies between when the cagemates were not soaked and when they were soaked (the aforementioned experiment). A two-way ANOVA for mixed design with a between-subjects factor of group (Distress; the cagemate was soaked in water, No Distress; the cagemate was not soaked in water) and a within-subject factor of session (7) on the latency of door-opening behavior revealed main effects of group (*F*(1, 24) = 12.62, *p* = .002) and session (*F*(6, 144) = 11.12, *p* < .001), while the effect of group × session interaction was close to significant (*F*(6, 144) = 1.95, *p* = .077). This suggests that the learning of the door-opening behavior is prevented when cagemates are not soaked in water. Thus, the negative emotions of the cagemate in the aversive situation may be necessary for the learning of door-opening behavior in the helpers.

Next, to examine the interest in the soaked cagemate, we measured how long the helper vole stayed close to the soaked vole side in the ground area during the door-opening task. After dividing the ground area into near and far halves relative to the pool area, the staying time of the helper voles in the two areas was measured (Figure 1F). The data of three voles with incomplete video recordings were excluded. A two-way ANOVA for mixed design with a between-subjects factor of group (Distress, No Distress) and a within-subject factor of session (7) revealed a main effect of session (*F*(6, 126) = 2.37, *p* = .033). Neither a main effect of group (*F*(1, 21) = 0.90, *p* = .355) nor a group × session interaction (*F*(6, 126) = 1.53, *p* = .174) was observed. This suggests that the interest of the helper voles in the cagemates in the pool area is not influenced by whether they are soaked or not.

In addition, we measured the duration of huddling in the pairs of helper and soaked voles during the two-minutes interaction period after the door-opening as an indicator of social attachment (Figure 1G). Huddling was defined as touching part of each other’s trunks. A two-way ANOVA for mixed design with a between-subjects factor of group (Distress, No Distress) and a within-subject factor of session (7) on the duration of huddling revealed neither a main effect of group (*F*(1, 21) = 0.12, *p* = .731) nor a group × session interaction (*F*(6, 126) = 0.90, *p* = .496), while the effect of session was close to significant (*F*(6, 126) = 1.95, *p* = .077). This suggests that the social attachment of prairie voles after the door-opening behavior is not affected by whether their cagemates are soaked in water.

### No Effect of the Sex Combination of the Pair on Helping Behavior in Prairie Voles

We examined the effect of the sex combination of the pair on helping behavior in prairie voles. Seven pairs of all sex combinations (Figure 2A) were tested. The latency of the door-opening appears to be decreased over the seven sessions in both the same-sex pairs and the opposite-sex pairs (Figures 2B, 2C and Figure S1). A two-way ANOVA for mixed design with a between-subjects factor of the sex combinations and a within-subject factor of session (7) revealed no main effect of sex (*F*(3, 24) = 0.86, *p* = .477). There was a significant effect of session (*F*(6, 144) = 23.35, *p* < .001), while the effect of a sex × session interaction was close to statistical significance (*F*(18, 144) = 1.55, *p* = .080). This suggests that there is no difference in the learning of helping behavior. However, the opposite-sex pairs may be delayed in learning helping behaviors.

**Figure 2.**
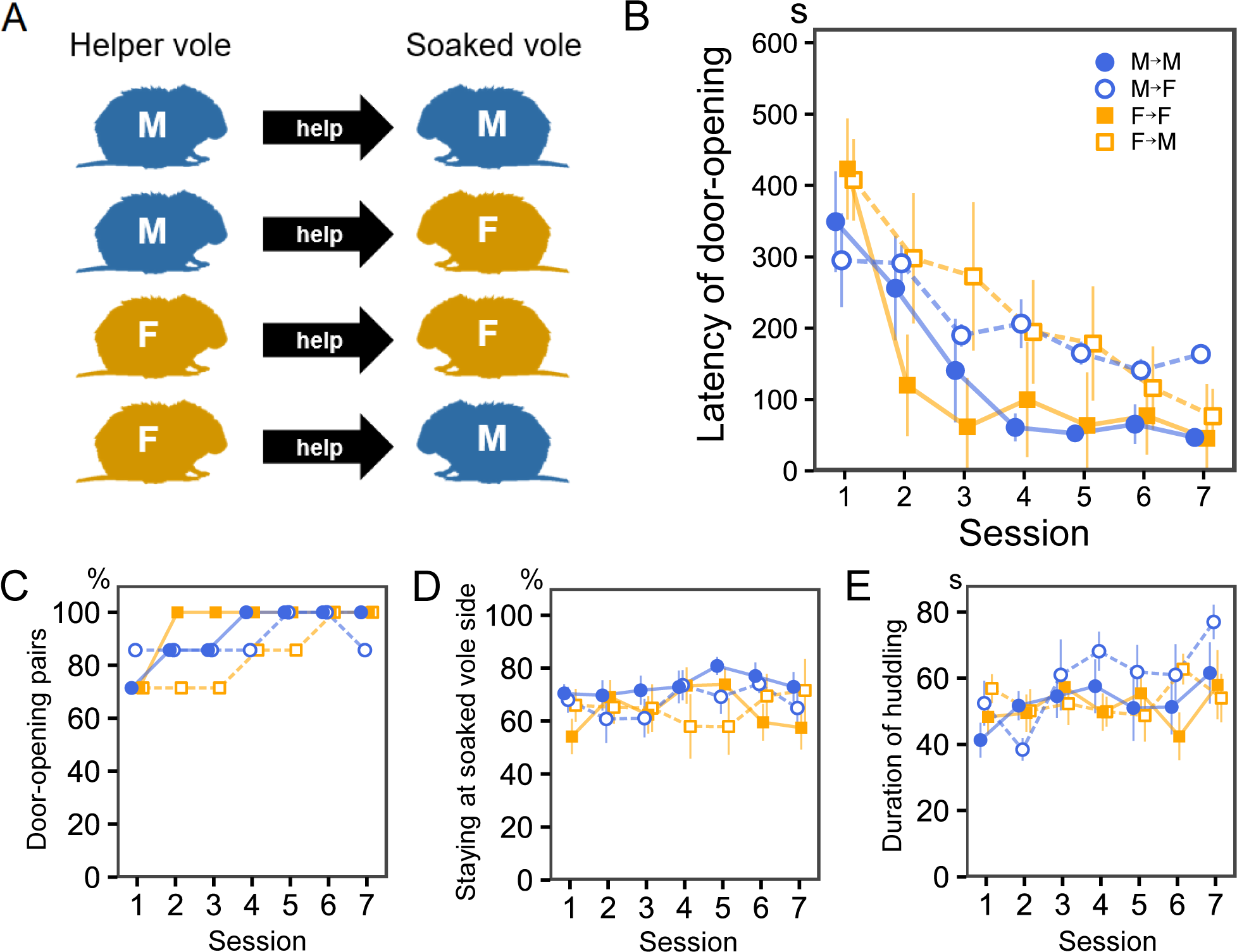
No differences in door-opening behavior were found between different sex combinations of pairs. (A) All sex combinations of the helper and soaked voles in this experiment. (B) The mean latency of door-opening in the helper voles of all sex combinations (n = 7 / group). (C) The percentage of door-opening pairs of all sex combinations. (D) The percentage of time that the helper of all sex combinations stayed on the soaked vole side. (E) The mean duration of huddling after opening the door in all sex combinations.

In addition, we assessed the percentage of time that the helper voles stayed on the soaked vole side during the door-opening task (Figure 2D). A two-way ANOVA for mixed design with a between-subjects factor of group and a within-subject factor of session revealed neither an effect of group (*F*(3, 24) = 0.79, *p* = .512), an effect of session (*F*(6, 144) = 0.39, *p* = .885), nor a group × session interaction (*F*(18, 144) = 0.99, *p* = .479). This suggests that there is no difference in interest of the helper voles in the soaked cagemates between the same-sex and opposite-sex pairs. In addition, we investigated the duration of huddling in the pairs of all sex combinations (Figure 2E). A two-way ANOVA on the duration of huddling revealed a main effect of session (*F*(6, 144) = 2.39, *p* = .031). Neither a main effect of group (*F*(3, 24) = 0.63, *p* = .601) nor a group × session interaction (*F*(18, 144) = 1.20, *p* = .266) was observed. This suggests that the sex combination of the pair has no effect on social attachment.

### Oxytocin Receptor Knockout Voles Demonstrate Less Helping Behavior

To investigate the impact of the elimination of oxytocin function on helping behavior, we examined the learning of door-opening behavior in oxytocin receptor knockout (OXTr -/-; OXTrKO) voles and compared it with that in wildtype voles (OXTr +/+; n = 14, the same-sex pairs in the aforementioned experiment). The OXTrKO voles were homozygous for the knocked out oxytocin receptor gene. The OXTrKO voles were paired with wildtype littermates (Figure 3A). All OXTrKO voles (n = 8) were assigned to be helpers. The OXTrKO and wildtype voles were same-sex pairs.

**Figure 3.**
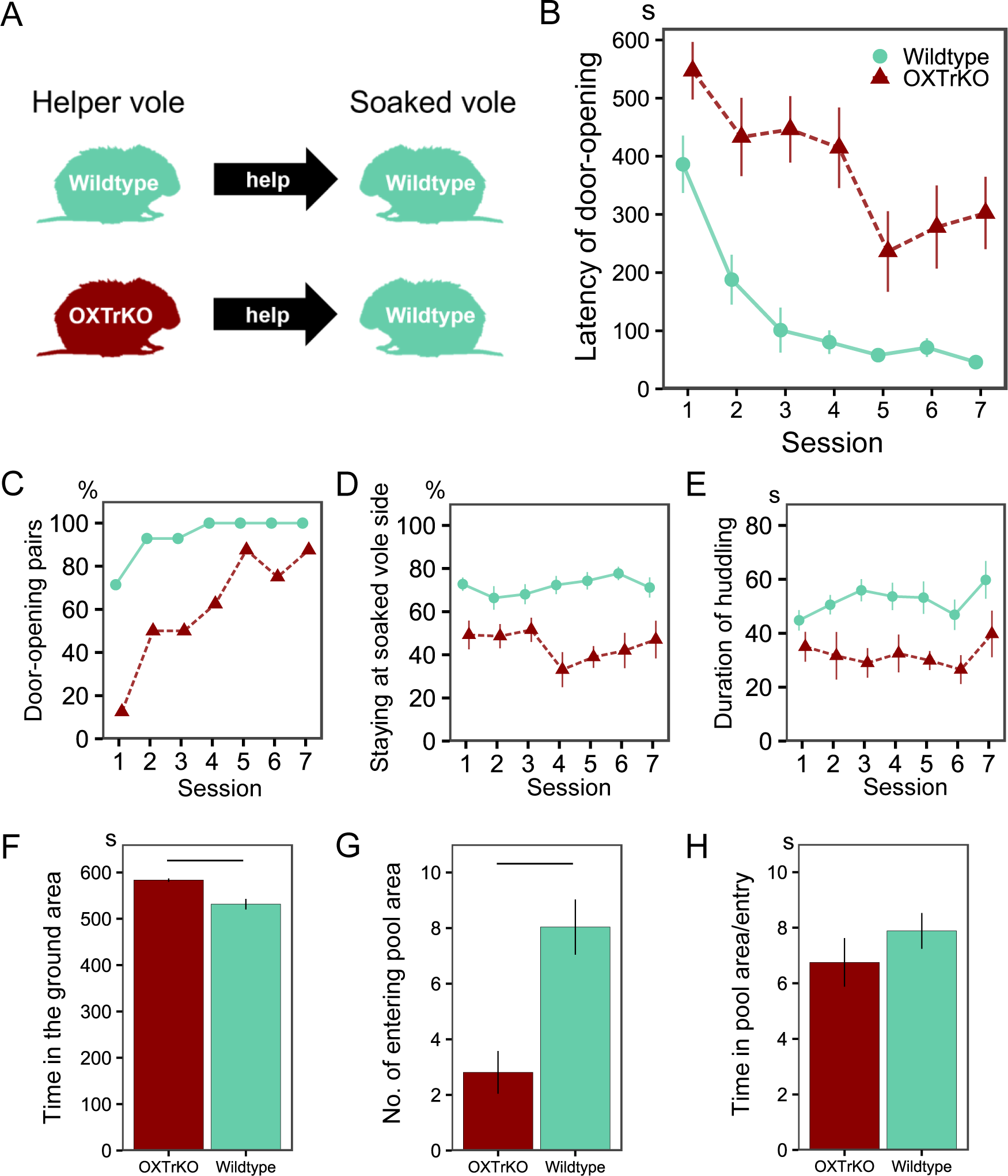
OXTrKO voles show less learning of helping behavior. (A) The comparison of the door-opening behavior between the wildtype and oxytocin receptor knockout (OXTrKO) helper voles. (B) The mean latency of door-opening in helper voles of the wildtype (n = 14) and OXTrKO voles (n = 8). (C) The percentage of door-opening pairs of the wildtype and OXTrKO voles. (D) The percentage of time that the wildtype and OXTrKO voles stayed on the soaked vole side. (E) The mean duration of huddling after opening the door in the wildtype and OXTrKO voles. (F) The time spent in the ground area in the OXTrKO and wildtype voles. The black line indicates a significant difference (*p* < .001). (G) The number of entries into the pool area in the OXTrKO and wildtype voles. (H) The time spent in the pool area per entry in the OXTrKO and wildtype voles.

The OXTrKO voles did not demonstrate a consistent decrease in the latency of door-opening behavior (Figure 3B). Two voles showed no consistent door-opening even after the fourth session (Figure 3C and Figure S3). A one-way ANOVA revealed a significant effect of session (*F*(6, 42) = 5.84, *p* < .001). However, Holm’s multiple comparisons revealed no significant difference in the latency of door-opening between all the sessions. The latency of door-opening was compared between the OXTrKO and wildtype voles. A two-way ANOVA for mixed design with a between-subjects factor of group (OXTrKO, Wildtype) and a within-subject factor of session (7) revealed main effects of group (*F*(1, 20) = 27.93, *p* < .001) and session (*F*(6, 120) = 18.84, *p* < .001), while the effect of a group × session interaction was close to significant (*F*(6, 120) = 2.01, *p* = .070).

Next, we compared the time the helper vole on the soaked vole side during the door-opening task in the OXTrKO voles (Figure 3D). A two-way ANOVA for mixed design revealed a main effect of group (*F*(1, 20) = 41.87, *p* < .001), but neither an effect of session (*F*(6, 120) = 0.57, *p* = .757) nor a group × session interaction (*F*(6, 120) = 1.55, *p* = .169). These results suggest that the OXTrKO voles demonstrate less helping behavior and show less interest in the soaked voles than the wildtype voles do. Additionally, we examined the duration of huddling in the OXTrKO voles (Figure 3E). A two-way ANOVA for mixed design on the duration of huddling revealed a main effect of group (*F*(1, 20) = 9.49, *p* = .006). Neither a main effect of session (*F*(6, 120) = 1.51, *p* = .181) nor a group × session interaction (*F*(6, 120) = 0.66, *p* = .685) was observed. This suggests that oxytocin affects social attachment in prairie voles.

To evaluate the water aversion of the OxtrKO voles, we compared the time spent in the ground and pool areas. During the habituation period, the OXTrKO voles stayed longer in the ground area (583.56±3.53 s, mean±SD) than the pool area, and the time spent in the ground area was significantly longer than the expected value (300 s, half of the habituation period, *t*(7) = 179.29, *p* < .001). We compared the water aversion between the OXTrKO and wildtype voles. A Welch’s t-test showed that the time spent in the ground area was longer in the OXTrKO voles than in the wildtype voles (Figure 3F; *t*(14.42) = 5.45, *p* < .001). The number of entries into the pool area was significantly higher in the wildtype voles than in the OXTrKO voles (Figure 3G; *t*(19.09) = 3.73, *p* < .001). No significant difference was found between the OXTrKO and wildtype voles in time spent in the pool area per entry (Figure 3H; *t*(13.68) = 0.98, *p* = .342). These results suggest that there is no difference in water aversion between the OXTrKO and wildtype voles.

## DISCUSSION

In this study, we investigated helping behavior in prairie voles using the door-opening paradigm. The results demonstrated that the helper voles quickly learned the door-opening behavior to free the soaked voles from water. As compared to when the cagemates were soaked, when they were not soaked in water the prairie voles did not immediately learn to open the door. This suggests that the cagemate being in a distressed situation is necessary for the learning of door-opening behavior. In addition, we found no sex difference in the latency of door-opening nor in the interest in the soaked vole. The OXTrKO voles demonstrated less learning of door-opening behavior and less interest in the soaked vole than the wildtype voles. This suggests that oxytocin is important for the emergence of helping behavior.

Helping behavior has been observed in rats when conspecifics are distressed situations, such as being trapped (Bartal et al., 2011) or soaked in water (Sato et al., 2015). It has been explained that helping behavior is learned through emotional contagion, a form of empathy, which involves sharing the distress of others (Bartal et al., 2011; Decety et al., 2016; Sato et al., 2015). In the present study, the helper vole could see the soaked vole through a transparent acrylic plate. Perceiving the conspecifics’ distress might drive the helping behavior in the helper voles. When the helper voles observed the soaked voles in distress, they might have shared the soaked voles’ distress through emotional contagion, and this might have motivated the helper voles to open the door (Sato et al., 2015).

When their cagemates were not soaked in water, the prairie voles learned less helping behavior. There is a claim that in rats, social interaction instead of empathy motivates door-opening behavior (Silberberg et al., 2014), although this has been rebutted (Sato et al., 2015). In this study, when the cagemate was not soaked in water, the learning of door-opening in the helper voles was not evident. This situation in which the cagemate was not distressed did not induce empathy in the helper vole, while the helper and soaked voles were still able to socially interact after the door was opened. This implies that empathy for distressed others, rather than social interaction, is important for helping behavior in prairie voles. Previous studies in rats have also shown that learning of helping behavior is suppressed when others are not in distressed situations (Bartal et al., 2011; Sato et al., 2015). Additionally, it was suggested that helping behaviors may occur even when the effects of social interaction are excluded (Cox and Reichel, 2019). These studies support the view that empathy for the distress of others may be more influential than social interaction as a trigger for door-opening behavior.

Oxytocin works as a hormone and as a neurotransmitter, which is involved in a variety of sociality, such as nurturing, familiarity, empathy, and social attachment (Bosch and Neumann, 2012; Decety et al., 2016; Insel, 2010; King et al., 2016; Rodrigues et al., 2009; Young and Wang, 2004). Many studies have shown that oxytocin and oxytocin receptors are involved in a variety of social behaviors in humans, non-human primates, and rodents (Amico et al., 2004; Anacker and Beery, 2013; Ferguson et al., 2000; Pobbe et al., 2012; Rich et al., 2014). Especially in mice, previous studies indicated that oxytocin gene knockout caused a lack of social recognition (Bielsky and Young, 2004; Ferguson et al., 2001), formation of social bonds (Liu and Wang, 2003; Young and Wang, 2004), and social rewards (Dölen et al., 2013; Hung et al., 2017). High sociality in prairie voles may be related to a higher density of oxytocin receptors in the brain as compared to close relative species (Olazábal and Young, 2006; Ross et al., 2009). A previous study reported that oxytocin receptor knockout voles demonstrated autism-like behavior such as a lack of interest in social novelty (Horie et al., 2019). In the present study, the decrease in the latency of the door-opening in the OXTrKO voles was slower than in the wildtype voles. The duration of huddling in the OXTrKO voles was also shorter than that in the wildtype voles. These findings suggest that blocking the function of oxytocin has a significant impact on helping behavior and social attachment. Nurturing and social play behavior in oxytocin knockout or oxytocin receptor knockout rodents as well as wildtype rodents is influenced by social context (Bredewold et al., 2014) and sexuality (Nishimori et al., 1996; Young III et al., 1996; Zimmermann-Peruzatto et al., 2017). Helping behavior in oxytocin knockout voles may also be affected by social context and sexual interaction. Future studies will examine the interaction between the oxytocin receptor knockout and these social elements.

In the present study, soaking in water was set as an aversive situation. In the preference test, the helper voles stayed much longer in the ground area than the pool area, even when the helper voles could go back and forth between the ground and pool areas. This suggests that prairie voles have an aversion to water and is consistent with previous studies that reported water aversion in rats (Morris, 1984; Sato et al., 2015). In the OXTrKO voles, the time spent in the pool area was shorter and the frequency of entering the pool area was lower than in the wildtype voles. However, there was no difference in the time spent in the pool area per entry between the OXTrKO and wildtype voles. The difference in the time spent in the pool area between the OXTrKO and wildtype voles may be explained by the difference in the number of times the voles entered the pool area instead of a difference in the degree of water aversion.

Oxytocin has been gaining attention for its effects on social cognition in recent years. Specifically, several studies have reported that intranasal administration of oxytocin can alleviate some symptoms of autism (Guastella et al., 2010; Hollander et al., 2003; Preti et al., 2014). The present study implies a relationship between helping behavior and oxytocin in prairie voles. Further studies will be needed to manifest the details, and they will shed light on the mechanisms of social cognition and related psychological disorders. Experiments in prairie voles will be beneficial for examining social cognitive functions.

## LIMITATION OF STUDY

All of the helper and soaked voles in the present study were littermates. Various factors of social context are involved in empathy (Bartal et al., 2014), such as familiarity and kinship. A previous study examining the emotional contagion of pain in mice suggests that the pain of familiar individuals is more contagious than that of strangers (Langford et al., 2006). Prairie voles exhibit allogrooming that functions as social buffering to relieve the fear of their partner. Allogrooming of a cagemate or a sibling is observed more frequently than that of a stranger (Burkett et al., 2016). The fact that the helper and soaked voles were littermates in the present study might have made the learning of the door-opening behavior easier than if they were strangers to each other. Rats display helping behavior not only toward cagemates but also toward strangers (Bartal et al., 2014, Yamagishi et al., 2019). It is unclear whether prairie voles exhibit helping behavior with strangers. The effects of familiarity and kinship on helping behavior in prairie voles should be investigated in the future.

## RESOURCE AVAILABILITY LEAD CONTACT

Further requests should be directed to and will be fulfilled by the Lead Contact: Nobuya Sato.

## MATERIALS AVAILABILITY

The study did not generate any unique reagent.

## DATE AND CODE AVAILABILITY

Data and code can be obtained from the corresponding author on request.

## ACKNOWLEDGMENTS

The authors thank Dr. Yoshiyuki Takahashi, Mr. Tomohiro Hayashi, and Ms. Shiori Michino for helpful discussion and animal breeding. This work was supported by JSPS KAKENHI Grant Numbers 16H01490 and 18K07357 and the Ministry of Education, Culture, Sports, Science and Technology (MEXT), JAPAN, for the Strategic Research Foundation at Private Universities (2015-2019; Project number S1511032) grant to the Center for Applied Psychological Science (CAPS), Kwansei Gakuin University.

## AUTHOR CONTRIBUTIONS

KK, AY, and NS designed the experiments. KK conducted the experiments and analyzed the data and wrote this manuscript with help from NS. AY created the experimental apparatus. KN and KH provided the animals. KN and NS reviewed and commented on this manuscript. NS acquired the funding.

## DECLARATION OF INTERESTS

The authors declare no competing interests.

## STAR*Methods

### Subjects

We used 80 experimentally naïve wildtype (52 males and 28 females) and 16 naïve oxytocin receptors knockout (OXTrKO, 8 males and 8 females) prairie voles (*Microtus ochrogaster*) maintained in a vivarium at Kwansei Gakuin University. The wildtype voles were captured in Illinois, USA. After that, the wildtype and the OXTrKO voles maintained at Tohoku University were delivered to Kwansei Gakuin University. The OXTrKO voles were generated at Tohoku University (v4 strain in Horie et al., 2019) and bred at Kwansei Gakuin University. At the beginning of the experiments, the mean age of the prairie voles was 21.1 weeks (range: 10–62 weeks) and the mean weight was 38.3 g (range: 25–63 g). They were housed in plastic home cages (320 × 212 × 130 mm, or 425 × 265 × 155 mm) with paper-chip bedding. The average number of prairie voles per cage was 3.9 (range: 2–6). Prairie voles in the same cage were littermates and were individually marked by ear punches. Helper and soaked roles were randomly assigned to individuals in the same cage. All animals were allowed free access to food (Labo MR Stock, NOSAN, Japan) and water. The vivarium was maintained at a constant temperature of 22 °C and 60% humidity. The light-dark cycle was set to 12:12 h with lights on at 9:00 am. This experiment was approved by the Animal Experimentation Committee of Kwansei Gakuin University (2018–44, 2019–06).

### Genotyping

OXTrKO voles were created in Horie et al. (2019). Genotyping to discriminate between wildtype (OXTr+/+) and OXTrKO (OXTr-/-) voles was done using the method of Horie et al. (2019). To summarize, we collected samples from prairie voles’ ear tissues and used them for a polymerase chain reaction. In the polymerase chain reaction, the forward polymer was 5’-AGA TCA GTG CCC GGG GGG TGC CC-3’ and the reverse polymer was 5′-TCG AGC GAC ATA AGC AGC AG-3’. We used prairie voles that were homozygous for the knocked out oxytocin receptor gene in the experiment.

### Experimental setup

The experimental apparatus was similar to that used in Sato et al. (2015). The size of the experimental box was smaller (240 × 120 × 210 mm) and was made of gray polyvinyl chloride boards (5 mm in thickness). It consisted of ground and pool areas separated by a transparent acrylic plate (5 mm in thickness). During the experiment, the pool area contained water with a depth of 25 mm to create an aversive situation for the prairie voles. The transparent acrylic plate had a circular hole (51 mm in diameter). On the ground area side, the hole was covered with a circular door (60 mm in diameter, 3 mm in thickness). The door completely separated the ground area from the pool area. A helper vole in the ground area opened the door, allowing a soaked vole to escape from the pool area to the ground area. The difficulty of opening the circular door was set to a constant level by holding it between two transparent acrylic fragments (Figure 1A). One pushed up the circular door with springs from below and the other was attached to the transparent acrylic plate. To prevent soaked voles from interfering with the circular door, a thin transparent sheet was put on the surface of the pool area side of the transparent acrylic plate. The transparent sheet had three small holes in the area that overlapped the circular door. After the start of the experiment, two thin transparent plates (150 × 150 mm, 5 mm in thickness) were placed on top of the experimental apparatus to prevent the voles from escaping out of it.

### Habituation

Before the experiment, we habituated the helper voles to the experimental apparatus. The habituation was performed for 10 minutes per trial for two days for each helper vole. During the habituation, the door was detached from the dividing plate and was laid on the floor of the ground area. The helper voles could move freely between the ground and pool areas during the habituation. The pool area was filled with water.

### Task procedure

Before the door-opening task, the experimental apparatus was cleaned with a 20% alcohol solution, the circular door was placed on the plate, and water was poured into the pool area to a depth of 25 mm. Immediately after the soaked vole was placed in the pool area, the helper vole was placed in the ground area. We measured the latency of the door-opening from the placement of the helper vole in the ground area. A trial of the task was carried out for a maximum of 10 minutes. If the helper vole opened the door in 10 minutes, we pulled out the sheet attached on the pool side of the plate to allow the soaked vole to escape to the ground area and the two voles to interact with each other. The duration of the interaction was two minutes. If the helper vole did not open the door within 10 minutes, the experimenter slightly opened the door to the right side and continued the trial for five more minutes. When the door-opening behavior was observed during those five minutes, the transparent sheet was removed to allow for interaction. If the helper vole did not open the door in the extra five minutes, the experimenter opened the circular door completely and removed the sheet to allow the interaction. This door-opening task was carried out in one trial per day, for a total of seven days.

### Data analysis

We analyzed the latency of door-opening and the time that the helper vole stayed on the soaked vole side during the first 10 minutes of the task. The soaked vole side was defined as the pool area side of the ground area divided in half. If the helper vole did not open the door for 10 minutes, the latency of door-opening of the trial was recorded as 600 seconds. We also measured whether the helper and soaked voles displayed huddling in two minutes of the interaction period after the door-opening. Huddling was defined as touching parts of each other’s trunks. During the habituation of the helper voles, we measured the time that the helper voles stayed in the ground and pool areas. The experiment was recorded by a video camera (HDR-CX590, Sony) mounted above the experimental apparatus. RStudio (version 1.1.453) was used for all statistical analyses.

Supplemental Video 1. An example scene of the distress condition

Supplemental Video 2. An example scene of the no distress condition

## Notes

### Competing Interest Statement

The authors have declared no competing interest.

## REFERENCES

Amico, J. A., Mantella, R. C., Vollmer, R. R., and Li, X. (2004). Anxiety and stress responses in female oxytocin deficient mice. Journal of Neuroendocrinology, 16, 319–324. doi: 10.1111/j.0953-8194.2004.01161.x

Anacker, A. M. J., and Beery, A. K. (2013). Life in groups: The roles of oxytocin in mammalian sociality. Frontiers in Behavioral Neuroscience, 7, 185. doi: 10.3389/fnbeh.2013.00185

Aragona, B. J., and Wang, Z. (2004). The prairie vole (Microtus ochrogaster): An animal model for behavioral neuroendocrine research on pair bonding. ILAR Journal, 45, 35–45. doi:10.1093/ilar.45.1.35

Atsak, P., Orre, M., Bakker, P., Cerliani, L., Roozendaal, B., Gazzola, V., Moita, M., and Keysers, C. (2011). Experience modulates vicarious freezing in rats: A model for empathy. PLoS One, 6, e21855. doi: 10.1371/journal.pone.0021855

Bartal, I. B. A., Decety, J., and Mason, P. (2011). Empathy and pro-social behavior in rats. Science, 334, 1424–1427. doi: 10.1126/science.1210789

Bartal, I. B. A., Rodgers, D. A., Sarria, M. S. B., Decety, J., and Mason, P. (2014). Pro-social behavior in rats is modulated by social experience. Elife, 3, e01385. doi: 10.7554/eLife.01385

Bartal, I. B. A., Shan, H., Molasky, N. M. R., Murray, T. M., Williams, J. Z., Decety, J., and Mason, P. (2016). Anxiolytic treatment impairs helping behavior in rats. Frontiers in Psychology, 7, 850. doi: 10.3389/fpsyg.2016.00850

Bartz, J. A., Zaki, J., Bolger, N., Hollander, E., Ludwig, N. N., Kolevzon, A., and Ochsner, K. N. (2010). Oxytocin selectively improves empathic accuracy. Psychological Science, 21, 1426–1428. doi: 10.1177/0956797610383439

Bielsky, I. F., and Young, L. J. (2004). Oxytocin, vasopressin, and social recognition in mammals. Peptides, 25, 1565–1574. doi: 10.1016/j.peptides.2004.05.019

Bosch, O. J., and Neumann, I. D. (2012). Both oxytocin and vasopressin are mediators of maternal care and aggression in rodents: From central release to sites of action. Hormones and Behavior, 61, 293–303. doi: 10.1016/j.yhbeh.2011.11.002

Bredewold, R., Smith, C. J. W., Dumais, K. M., and Veenema, A. H. (2014). Sex-specific modulation of juvenile social play behavior by vasopressin and oxytocin depends on social context. Frontiers in Behavioral Neuroscience, 8, 216. doi: 10.3389/fnbeh.2014.00216

Burkett, J. P., Andari, E., Johnson, Z. V., Curry, D. C., de Waal, F. B., and Young, L. J. (2016). Oxytocin-dependent consolation behavior in rodents. Science, 351, 375–378. doi: 10.1126/science.aac4785

Campbell, M. W., and de Waal, F. B. M. (2011). Ingroup-outgroup bias in contagious yawning by chimpanzees supports link to empathy. PLoS One, 6, e18283. doi: 10.1371/journal.pone.0018283

Carter, C. S., and Getz, L. L. (1993). Monogamy and the prairie vole. Scientific American, 268, 100–106. doi: 10.1038/scientificamerican0693-100

Cox, S. S., and Reichel, C. M. (2019). Rats display empathic behavior independent of the opportunity for social interaction. Neuropsychopharmacology, 45, 1097–1104. doi: 10.1038/s41386-019-0572-8

Cronin, K. A. (2012). Prosocial behaviour in animals: The influence of social relationships, communication and rewards. Animal Behaviour, 84, 1085–1093. doi: 10.1016/j.anbehav.2012.08.00

Decety, J., Bartal, I. B. A., Uzefovsky, F., and Knafo-Noam, A. (2016). Empathy as a driver of prosocial behaviour: Highly conserved neurobehavioural mechanisms across species. Philosophical Transactions of the Royal Society B: Biological Science, 371. 20150077. doi: 10.1098/rstb.2015.0077

de Waal, F. B. M., and Preston, S. D. (2017). Mammalian empathy: Behavioural manifestations and neural basis. Nature Reviews Neuroscience, 18, 498–509. doi: 10.1038/nrn.2017.72

Dölen, G., Darvishzadeh, A., Huang, K. W., and Malenka, R. C. (2013). Social reward requires coordinated activity of nucleus accumbens oxytocin and serotonin. Nature, 501, 179–184. doi: 10.1038/nature12518

Ferguson, J. N., Aldag, J. M., Insel, T. R., and Young, L. J. (2001). Oxytocin in the medial amygdala is essential for social recognition in the mouse. Journal of Neuroscience, 21, 8278–8285. doi: 10.1523/JNEUROSCI.21-20-08278.2001

Ferguson, J. N., Young, L. J., Hearn, E. F., Matzuk, M. M., Insel, T. R., and Winslow, J. T. (2000). Social amnesia in mice lacking the oxytocin gene. Nature Genetics, 25, 284–288. doi: 10.1038/77040

Gallup, A. C., Swartwood, L., Militello, J., and Sackett, S. (2015). Experimental evidence of contagious yawning in budgerigars (Melopsittacus undulatus). Animal Cognition, 18, 1051–1058. doi: 10.1007/s10071-015-0873-1

Getz, L. L., and Carter, C. S. (1996). Prairie-vole partnerships: This rodent forms social groups that appear to have evolved as an adaptation for living in a low-food habitat. American Scientist, 84, 56–62. from www.jstor.org/stable/29775598

Getz, L. L., and Hofmann, J. E. (1986). Social organization in free-living prairie voles (Microtus ochrogaster). Behavioral Ecology and Sociobiology, 18, 275–282. doi: 10.1007/BF00300004

Guastella, A. J., Einfeld, S. L., Gray, K. M., Rinehart, N. J., Tonge, B. J., Lambert, T. J., and Hickie, I. B. (2010). Intranasal oxytocin improves emotion recognition for youth with autism spectrum disorders. Biological Psychiatry, 67, 692–694. doi: 10.1016/j.biopsych.2009.09.020

Hollander, E., Novotny, S., Hanratty, M., Yaffe, R., DeCaria, C. M., Aronowitz, B. R., and Mosovich, S. (2003). Oxytocin infusion reduces repetitive behaviors in adults with autistic and Asperger’s disorders. Neuropsychopharmacology, 28, 193–198. doi: 10.1038/sj.npp.1300021

Horie, K., Inoue, K., Suzuki, S., Adachi, S., Yada, S., Hirayama, T., Hidema, S., Young, L. J., and Nishimori, K. (2019). Oxytocin receptor knockout prairie voles generated by CRISPR/Cas9 editing show reduced preference for social novelty and exaggerated repetitive behaviors. Hormones and Behavior, 111, 60–69. doi: 10.1016/j.yhbeh.2018.10.011

Hung, L. W., Neuner, S., Polepalli, J. S., Beier, K. T., Wright, M., Walsh, J. J., Lewis, E. M., Luo, L., Deisseroth, K., Dölen, G., and Malenka, R. C. (2017). Gating of social reward by oxytocin in the ventral tegmental area. Science, 357, 1406–1411. doi: 10.1126/science.aan4994

Insel, T. R. (2010). The challenge of translation in social neuroscience: A review of oxytocin, vasopressin, and affiliative behavior. Neuron, 65, 768–779. doi: 10.1016/j.neuron.2010.03.005

King, L. B., Walum, H., Inoue, K., Eyrich, N. W., and Young, L. J. (2016). Variation in the oxytocin receptor gene predicts brain region–specific expression and social attachment. Biological Psychiatry, 80, 160–169. doi: 10.1016/j.biopsych.2015.12.008

Koski, S. E., and Sterck, E. H. M. (2010). Empathic chimpanzees: A proposal of the levels of emotional and cognitive processing in chimpanzee empathy. European Journal of Developmental Psychology, 7, 38–66. doi: 10.1080/17405620902986991

Kuczaj, S. A., Frick, E. E., Jones, B. L., Lea, J. S. E., Beecham, D., and Schnöller, F. (2015). Underwater observations of dolphin reactions to a distressed conspecific. Learning & Behavior, 43, 289–300. doi: 10.3758/s13420-015-0179-9

Langford, D. J., Crager, S. E., Shehzad, Z., Smith, S. B., Sotocinal, S. G., Levenstadt, J. S., Chanda, M. L., Levitin, D. J., and Mogil, J. S. (2006). Social modulation of pain as evidence for empathy in mice. Science, 312, 1965–1967. doi: 10.1126/science.1128322

Liu, Y., and Wang, Z. X. (2003). Nucleus accumbens oxytocin and dopamine interact to regulate pair bond formation in female prairie voles. Neuroscience, 121, 537–544. doi: 10.1523/JNEUROSCI.21-20-08278.2001

Marlin, B. J., and Froemke, R. C. (2017). Oxytocin modulation of neural circuits for social behavior. Developmental Neurobiology, 77, 169–189. doi: 10.1002/dneu.22452

Meyza, K. Z., Bartal, I. B. A., Monfils, M. H., Panksepp, J. B., and Knapska, E. (2017). The roots of empathy: Through the lens of rodent models. Neuroscience and Biobehavioral Reviews, 76, 216–234. doi: 10.1016/j.neubiorev.2016.10.028.

Morris, R. (1984). Developments of a water-maze procedure for studying spatial learning in the rat. Journal of Neuroscience Methods, 11, 47–60. doi: 10.1016/0165-0270(84)90007-4

Nishimori, K., Young, L. J., Guo, Q., Wang, Z., Insel, T. R., and Matzuk, M. M. (1996). Oxytocin is required for nursing but is not essential for parturition or reproductive behavior. Proceedings of the National Academy of Sciences, 93, 11699–11704. doi: 10.1073/pnas.93.21.11699

Olazábal, D. E., and Young, L. J. (2006). Species and individual differences in juvenile female alloparental care are associated with oxytocin receptor density in the striatum and the lateral septum. Hormones and Behavior, 49, 681–687. doi: 10.1016/j.yhbeh.2005.12.010

Palagi, E., Nicotra, V., and Cordoni, G. (2015). Rapid mimicry and emotional contagion in domestic dogs. Royal Society Open Science, 2, 150505. doi: 10.1098/rsos.150505

Pisansky, M. T., Hanson, L. R., Gottesman, I. I., and Gewirtz, J. C. (2017). Oxytocin enhances observational fear in mice. Nature Communications, 8, 2102. doi: 10.1038/s41467-017-02279-5

Pobbe, R. L. H., Pearson, B. L., Defensor, E. B., Bolivar, V. J., Young, W. S., Lee, H. J., Blanchard, D. C., and Blanchard, R. J. (2012). Oxytocin receptor knockout mice display deficits in the expression of autism-related behaviors. Hormones and Behavior, 61, 436–444. doi: 10.1016/j.yhbeh.2011.10.010

Preston, S.D., and de Waal, F.B.M. (2002). Empathy: Its ultimate and proximate bases. Behavioral Brain Science, 25, 1–20. doi: 10.1017/S0140525X02000018

Preti, A., Melis, M., Siddi, S., Vellante, M., Doneddu, G., and Fadda, R. (2014). Oxytocin and autism: A systematic review of randomized controlled trials. Journal of Child and Adolescent Psychopharmacology, 24, 54–68. doi: 10.1089/cap.2013.0040

Pruetz, J. D. (2011). Targeted helping by a wild adolescent male chimpanzee (Pan troglodytes verus): Evidence for empathy? Journal of Ethology, 29, 365–368. doi: 10.1007/s10164-010-0244-y

Rich, M. E., DeCárdenas, E. J., Lee, H. J., and Caldwell, H. K. (2014). Impairments in the initiation of maternal behavior in oxytocin receptor knockout mice. PLoS One, 9, e98839. doi: 10.1371/journal.pone.0098839

Rodrigues, S. M., Saslow, L. R., Garcia, N., John, O. P., and Keltner, D. (2009). Oxytocin receptor genetic variation relates to empathy and stress reactivity in humans. Proceedings of the National Academy of Sciences, 106, 21437–21441. doi: 10.1073/pnas.0909579106

Rogers-Carter, M. M., Varela, J. A., Gribbons, K. B., Pierce, A. F., McGoey, M. T., Ritchey, M., and Christianson, J. P. (2018). Insular cortex mediates approach and avoidance responses to social affective stimuli. Nature Neuroscience, 21, 404–414. doi: 10.1038/s41593-018-0071-y

Romero, T., Konno, A., and Hasegawa, T. (2013). Familiarity bias and physiological responses in contagious yawning by dogs support link to empathy. PLoS One, 8, e71365. doi: 10.1371/journal.pone.0071365

Ross, H. E., Freeman, S. M., Spiegel, L. L., Ren, X., Terwilliger, E. F., and Young, L. J. (2009). Variation in oxytocin receptor density in the nucleus accumbens has differential effects on affiliative behaviors in monogamous and polygamous voles. Journal of Neuroscience, 29, 1312–1318. doi: 10.1523/JNEUROSCI.5039-08.2009

Ross, H. E., and Young, L. J. (2009). Oxytocin and the neural mechanisms regulating social cognition and affiliative behavior. Frontiers in Neuroendocrinology, 30, 534–547. doi: 10.1016/j.yfrne.2009.05.004

Sato, N., Tan, L., Tate, K., and Okada, M. (2015). Rats demonstrate helping behavior toward a soaked conspecific. Animal Cognition, 18, 1039–1047. doi: 10.1007/s10071-015-0906-9

Schulte, B. A. (2000). Social structure and helping behavior in captive elephants. Zoo Biology, 19, 447–459. doi: 10.1002/1098-2361(2000)19:5<447::AID-ZOO12>3.0.CO;2-%23

Silberberg, A., Allouch, C., Sandfort, S., Kearns, D., Karpel, H., and Slotnick, B. (2014). Desire for social contact, not empathy, may explain “rescue” behavior in rats. Animal Cognition, 17, 609–618. doi: 10.1007/s10071-013-0692-1

Stetzik, L. A., Sullivan, A. W., Patisaul, H. B., and Cushing, B. S. (2018). Novel unconditioned prosocial behavior in prairie voles (Microtus ochrogaster) as a model for empathy. BMC Research Notes, 11, 1–6. doi: 10.1186/s13104-018-3934-0

Tabbaa, M., Paedae, B., Liu, Y., and Wang, Z. (2011). Neuropeptide regulation of social attachment: The prairie vole model. Comprehensive Physiology, 7, 81–104. doi: 10.1002/cphy.c150055

Wardwell, J., Watanasriyakul, W. T., Normann, M. C., Akinbo, O. I., McNeal, N., Ciosek, S., Cox, M., Holzapfel, N., Sujet, S., and Grippo, A. J. (2020). Physiological and behavioral responses to observing a sibling experience a direct stressor in prairie voles. Stress, 23, 444–456. doi: 10.1080/10253890.2020.1724950

Winslow, J. T., and Insel, T. R. (2002). The social deficits of the oxytocin knockout mouse. Neuropeptides, 36, 221–229. doi: 10.1054/npep.2002.0909

Yamagishi, A., Lee, J., and Sato, N. (2020). Oxytocin in the anterior cingulate cortex is involved in helping behaviour. Behavioural Brain Research, 393, 112790. https://doi.org/10.1016/j.bbr.2020.112790.

Yamagishi, A., Okada, M., Masuda, M., and Sato, N. (2019). Oxytocin administration modulates rats’ helping behavior depending on social context. Neuroscience Research, 153, 56–61. doi: 10.1016/j.neures.2019.04.001

Yamamoto, S., Humle, T., and Tanaka, M. (2012). Chimpanzees’ flexible targeted helping based on an understanding of conspecifics’ goals. Proceedings of the National Academy of Sciences, 109, 3588–3592. doi: 10.1073/pnas.1108517109

Young, L. J., and Barrett, C. E. (2015). Can oxytocin treat autism? Science, 347, 825–826. doi: 10.1126/science.aaa8120

Young, L. J., and Wang, Z. (2004). The neurobiology of pair bonding. Nature Neuroscience, 7, 1048–1054. doi: 10.1038/nn1327

Young III, W. S., Shepard, E., Amico, J., Hennighausen, L., Wagner, K. U., Lamarca, M. E., McKinney, C., and Ginns, E. I. (1996). Deficiency in mouse oxytocin prevents milk ejection, but not fertility or parturition. Journal of Neuroendocrinology, 8, 847–853. doi: 10.1046/j.1365-2826.1996.05266.x

Zimmermann-Peruzatto, J. M., Lazzari, V. M., Agnes, G., Becker, R. O., de Moura, A. C., Guedes, R. P., Lucion, A. B., Almeida, S., and Giovenardi, M. (2017). The impact of oxytocin gene knockout on sexual behavior and gene expression related to neuroendocrine systems in the brain of female mice. Cellular and Molecular Neurobiology, 37, 803–815. doi: 10.1007/s10571-016-0419-3

