## Supplemental Figures for "Helping Behavior in Prairie Voles: A Model of Empathy and the Importance of Oxytocin"

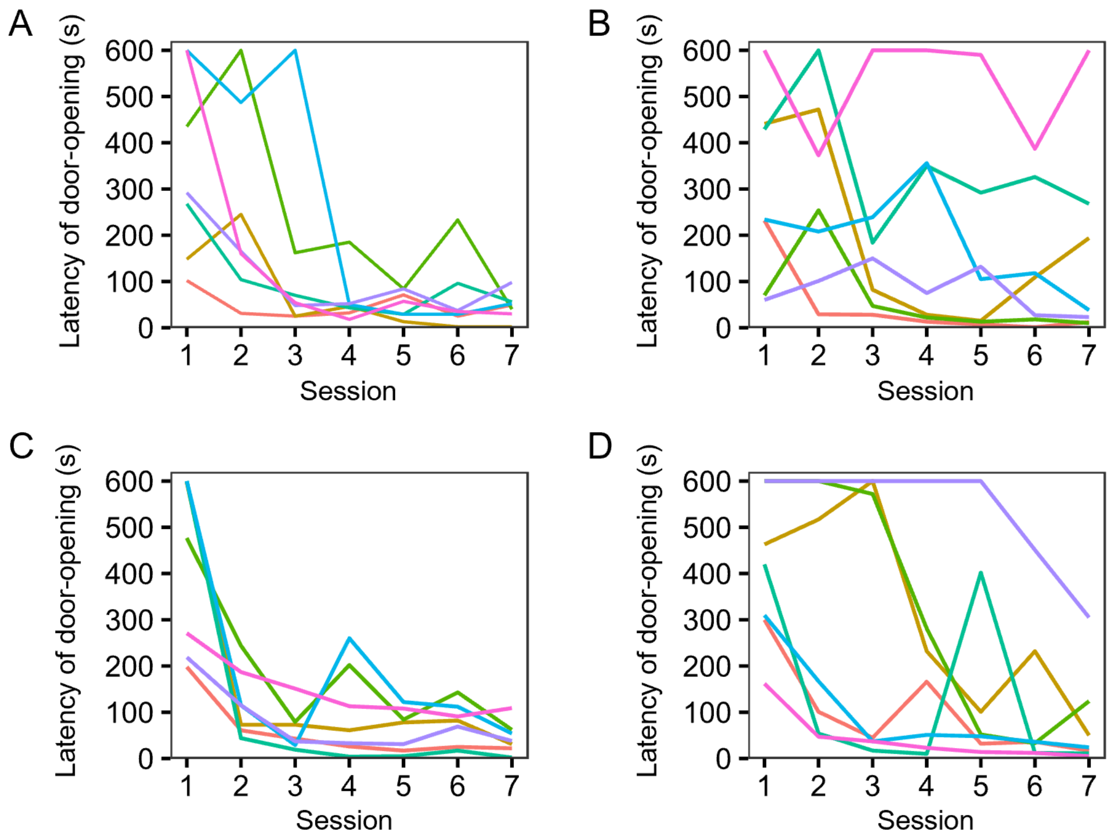


**Figure S1. Individual data of the helper voles in all sex combinations**

Individual latency of door-opening in all sex combinations of pairs: a male helping another male (A), a male helping a female (B), a female helping another female (C), a female helping a male (D).

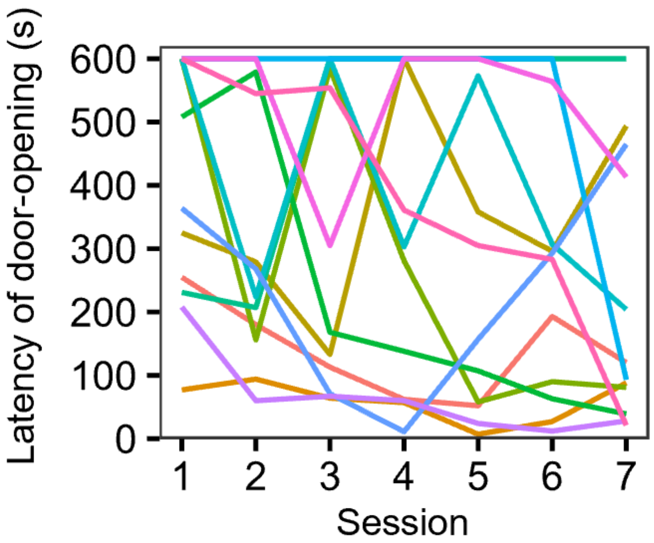


**Figure S2. Individual data of the helper voles when the cagemate was not soaked in water**

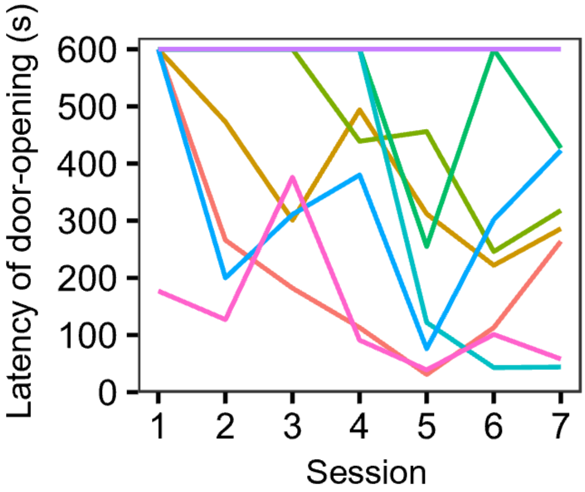


**Figure S3. Individual data of the helpers of the OXTrKO voles**
